# Reliable and Efficient Agentic Protein Design with Toolset Harnessing

**DOI:** 10.64898/2026.10.19.759896

**Authors:** Jasper Butcher, Jie Chen, Andrew Hunt, Cyrus M. Haas, Wenzhao Dai, Shajesh Sharma, Raktim Mitra, Saman Salike, Jeremy Koob, Aiko Muraishi, Zhe Li, Harley Pyles, Michael D. Tyka, Magnus S. Bauer, Rafi I. Brent, Anant Mittal, Neil P. King, David Baker

## Abstract

Computational protein design workflows require the orchestration of multiple tools, data formats, and analysis steps that are often fragmented. Here, we introduce Chaperone, a lightweight and customizable framework that enables seamless integration of RFdiffusion3 (RFD3) [1] with downstream sequence design, structure prediction, and evaluation tools, including ProteinMPNN [2], AlphaFold3 (AF3) [3], and more than 50 other protein design tools. By combining a programmable agent-friendly interface with parallelized execution on high-performance computing systems, Chaperone as a harness enables rapid iteration and scalable exploration of complex protein design workflows. Across an evaluation of nine biomolecular design tasks, we find using Chaperone reduces total token usage relative to baseline agents by 86% and GPU costs by 64%, while increasing the overall computational design pass rate. We demonstrate Chaperone’s utility for large-scale design-space exploration through a two-week end-to-end small-molecule binder design campaign, yielding low nanomolar binders to cortisol zero shot. We further demonstrate its versatility across distinct design challenges, including the design of a symmetric 12-subunit self-assembly nanoparticle and a linker that mediates interactions between C2- and D3-symmetric protein complexes to form crystalline assemblies. Together, these results demonstrate how agent harnesses built around standardized tool interfaces can substantially improve the efficiency and success of complex, long-horizon agentic design workflows in biology. We make Chaperone freely available to the community on GitHub: https://github.com/RosettaCommons/chaperone.

## 1 Introduction

Protein design pipelines have evolved rapidly in the recent years [2, 4–8]. The sprawling open-source ecosystem has spurred the development of a wide variety of deep learning-based generative methods [1,5,6,9], prediction oracles [3, 10–12], as well as physics-based scoring methods [13, 14]. This growing ecosystem, coupled with the increasing complexity of design tasks, often necessitates the integration of multiple such tools. Workflows are therefore often convoluted, requiring users to connect several independently developed tools into multi-step pipelines without standardized data formatting or APIs. Furthermore, users often rely on bespoke computational environments that require manual installation and scripting to execute these tools on their own systems. Collectively, these challenges create energy barriers for researchers seeking to explore novel design hypotheses through combinations of various tools. Several recent studies have demonstrated the importance of complex, multi-stage design strategies [15–19]. Widely adopted systems such as BindCraft [8] integrate multiple protein design tools into accessible, end-to-end pipelines. HalluDesign [16] and Protein Hunter [17] demonstrate the benefits of iterative design, improving computational outcomes through repeated cycles of generation, evaluation, and refinement. However, these systems are primarily built around predefined workflows, and incorporating alternative models, custom evaluation metrics, or new optimization strategies often requires modifying their underlying implementations.

We set out to address these challenges by developing a customizable and lightweight abstraction layer that integrates a broad range of protein design tools under one umbrella, with the goal of enabling seamless interoperability through a direct DataFrame-based framework. We hypothesized that such an interface would extend into a natural harness for AI agents to assist researchers in pursuing challenging protein design tasks, decreasing the barrier for exploration, and accelerating research beyond conventional timelines. We focused in particular on design workflows involving the powerful recently released RFD3 generative protein design program, widely used for enzyme, binder and conformational switch design.

## 2 The Chaperone framework

By interconnecting a broad set of design tools within a unified framework, Chaperone enables researchers to systematically explore complex design strategies, customize computational pipelines, and execute experiments in parallel across a diverse range of targets and tasks.

Previous work has addressed multi-tool orchestration challenges through several complementary approaches. BioPipelines [20] and ProteinDJ [21] facilitate multi-tool workflow construction and execution on high-performance computing systems, while ProtFlow [22] provides a DataFrame-based interface for managing candidate designs and their associated metrics. Proto [23] approaches biological design through programmable, constraint-based optimization, whereas ProteinMCP [24] and T-REX [25] demonstrate the potential of agent-driven tool orchestration and adaptive design campaigns. Building on these efforts, we sought to develop a maximally lightweight framework that simplifies the integration of heterogeneous protein design tools, motivated in particular by the broad range of design tasks enabled by RFdiffusion3 (RFD3). Applications ranging from enzyme scaffolding and small-molecule binder design to RNA-binding protein design, symmetric assemblies, and nanoparticles each require different combinations of computational tools, evaluation metrics, and design strategies. We therefore aimed to provide a common interface through which such workflows could be readily constructed, modified, and scaled without rewriting the underlying orchestration logic.

### Blocks and the shared DataFrame

We observed that most biomolecular design tools perform one of two operations: (i) generating new candidates from existing structures, or (ii) annotating existing candidates with additional metrics. Both can be represented through a common DataFrame schema, in which each row is one design and carries its structural history and metrics. We term each operation a *block*, a lightweight Python class that simply takes as input a DataFrame and returns an extended DataFrame (Fig. 1**a**). A block appends its output structure to each row’s history and adds its metrics alongside those already present. When a tool produces several candidates from one input, the parent row is duplicated for each output. Each row holds an ordered list of structure records (the producing step, file path and residue-level annotations) and a JSON-encoded collection of metrics, stored in a structured columnar schema built on Polars [26]. Generation, prediction and scoring blocks all share this interface. This preserves design lineage and keeps a single source of truth throughout a campaign.

**Figure 1.**
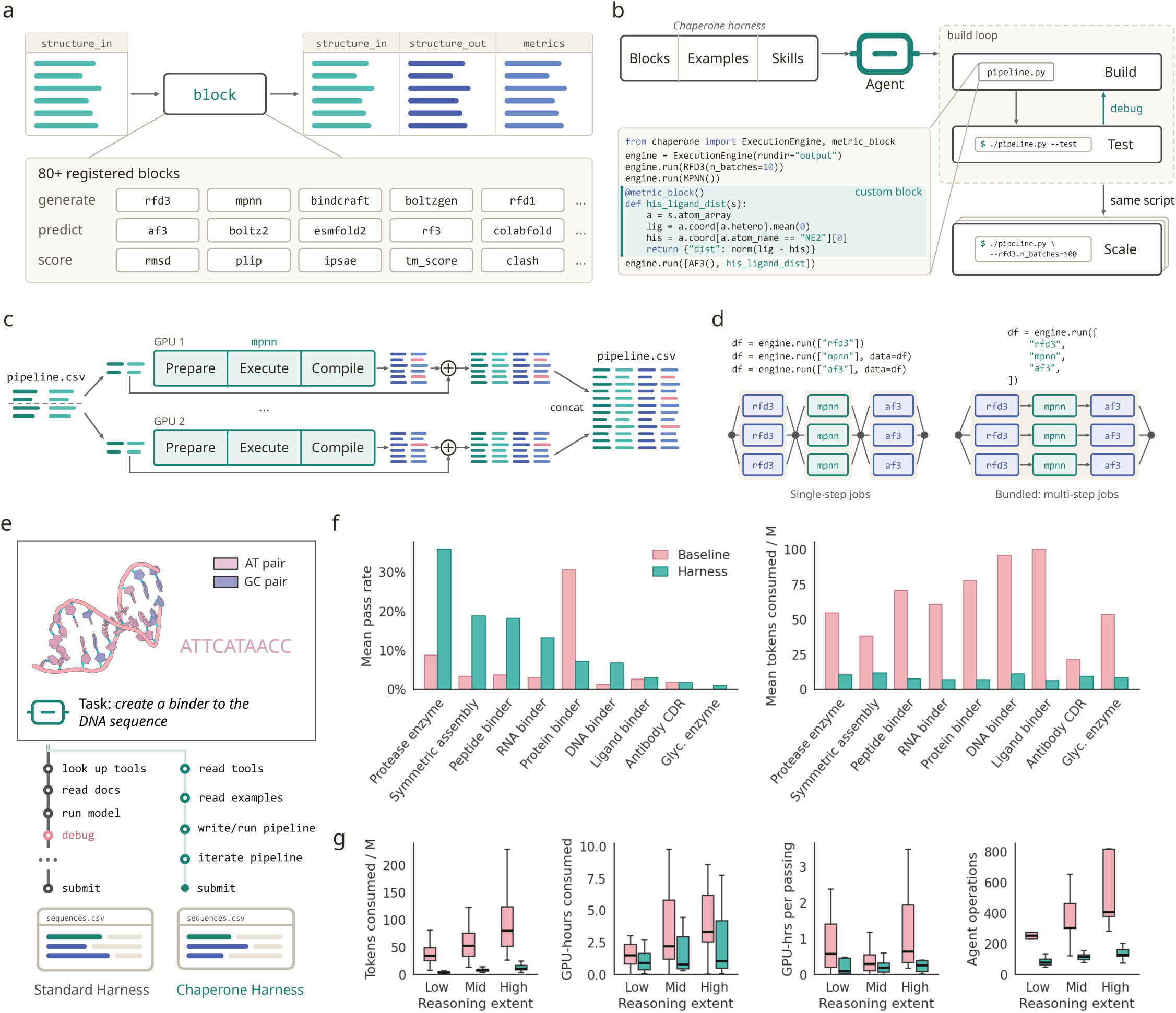
Overview and evaluation of the Chaperone harness for autonomous protein-design agents. **a**, The block API. Every block takes a DataFrame and returns a DataFrame. Each row is one design. A block adds its output structure to that row and can also add its own metrics. Generation, prediction and scoring blocks all follow this same interface, so any block can follow any other. The registry holds over 80 blocks. **b**, The Chaperone harness gives the agent three layers of context: blocks, examples and skills. The agent works in a build–test loop; writing one script and running it with the test override (which ensures every tool in the pipeline is run on lightweight settings.) This forms a verifiable loop for long workflows and ensures blocks work together end-to-end. Scaling can be done arbitrarily; changed by command-line overrides to any blocks executed. Inset: example pipeline.py combining common blocks (RFD3, MPNN, AF3) interleaved with arbitrary custom code written by the agent (ligand–histidine distance computed across all AF3 outputs.) **c**, The execution engine splits the input DataFrame into chunks and distributes them across GPUs. Each chunk runs the block’s three phases: prepare, execute and compile. The config supplied determines how inputs are prepared and how outputs are compiled. Each chunk’s compiled output is back-linked to that chunk’s input rows using path handles, which extends each row with the new structures and metrics. When all jobs finish, the chunks are concatenated into a single DataFrame. **d**, Pipeline steps can run separately or bundled. Left: in conventional execution, each run call executes one block, and its returned DataFrame is passed to the next call once outputs have been gathered. Right: in bundled execution, one call takes several blocks and runs them in sequence inside a single job. Jobs can be local processes, on SLURM/SGE or run on cloud computing. This can be combined with custom code in (B) to achieve fully customizable multi-step pipelines with maximal use of resources. **e**, Evaluation setup. Each agent receives a task and must submit 96 sequences, which held-out scorers judge. The agents do not see the scoring oracle. A standard Codex agent looks up tools, reads documentation, runs models and debugs. The Chaperone agent reads the harness context, then writes and iterates on a pipeline script. **f**, Mean pass rate (left) and mean tokens per run (right) for nine design tasks, without (baseline) and with the harness. Each bar averages nine runs (three replicates at each of three reasoning efforts). Compared against baseline, Chaperone is higher in six tasks, similar in two (ligand binder, antibody CDR) and lower for PD-L1. Token use is lower in all nine. **g**, Cost by reasoning effort for GPT-5.6 Sol run through Codex, with 27 runs per effort (3 attempts for nine tasks). Panels show tokens consumed, GPU-hours consumed, GPU-hours per passing design (runs with no passing design omitted) and agent operations per run. Boxes show median and interquartile range. Across all 81 runs, the harness raised the mean pass rate from 6.1% to 11.7% and cut tokens per run by 86% and GPU-hours per run by 64%.

### Executing a block (**Fig. 1c**)

Each block separates input preparation, execution and output compilation. The tool itself is treated as a black box, so its environment and input/output conventions stay unchanged. Preparation converts the input DataFrame into the files and parameters the tool expects; compilation collects the results and converts them back into the canonical DataFrame. For example, a ProteinMPNN [2] block prepares backbone inputs, runs sequence design, and attaches the generated sequences and metrics to their parent structures. The execution engine splits the input DataFrame into independent chunks and distributes them across jobs. Each chunk’s compiled output is merged with the original input DataFrame by tracing the lineage for each output to its source based on path handles. Chunks are concatenated when all jobs finish. Intermediate outputs are stored as Parquet files, and incomplete workflows can be resumed from previously completed steps or chunks. Local multiprocessing and HPC job submission run through a custom scheduler built around Submitit [27].

### Writing and scaling pipelines

A pipeline script can mix common registered blocks with arbitrary custom code to be executed during jobs, including custom operations (Fig. 1**b**). The syntax for writing pipelines is simple and follows a common pattern; an execution engine receives the configuration of the block to run and executes it with a supplied dataframe (Algorithm 1).

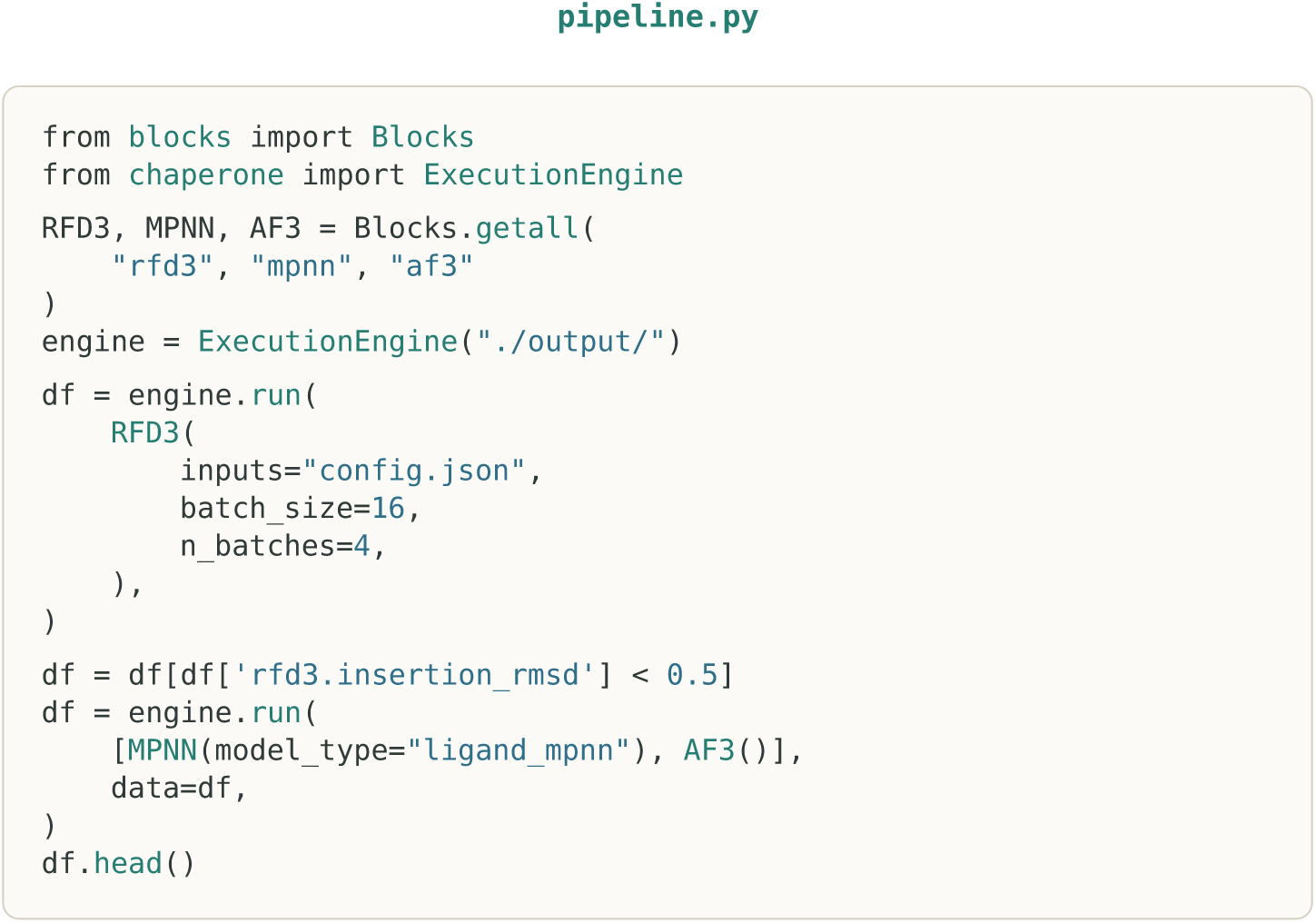

**Algorithm 1** Syntax for a basic RFD3-MPNN-AF3 enzyme design pipeline with a custom filtering step. Each call to engine.run applies one or more blocks to a dataframe with one row per design and returns it with new columns for the block outputs. Blocks are looked up by name, instantiated with their arguments, and passed to engine.run. RFD3 first generates backbones from a configuration file (four batches of 16). The dataframe is then filtered with ordinary indexing on the rfd3.insertion_rmsd metric (the custom filter, which can be any ordinary Python), and the surviving designs are passed to a single chained call that runs LigandMPNN sequence design (model_type=“ligand_mpnn”) followed by AF3 structure prediction.

**Agent harness** Given its programmatic nature, we find Chaperone is particularly well suited as an interface for agentic workflows. An agent harness refers to the set of native tools available to agents including compaction, and context management as well as toolsets, skills and MCPs. In Figure 1**b** we extend Chaperone as a harness for agents, providing the Chaperone library as the primary tool-use interface. The harness provides an agent four layers of context for writing and managing protein design workflows:

1. The abstract scripting layer: tool-independent ways to run design pipelines, interfaces for compute, the DataFrame-to-DataFrame API schema described above.
2. Documentation of blocks: since every block operates on a standardized schema, the output of each tool can be understood from the code alone, making the construction of the workflow devisable before any runtime execution.
3. Examples of integrated workflows across design tasks spanning both design tasks as well as tool-use edge-cases such as branching, cycling and templating.
4. Workflow orchestration in the form of agent skills, for example: effectively monitoring cluster jobs and resources, recovering crashed or incomplete outputs, scripting, debugging inputs, visualisation, setup and scripting.

Additional features of Chaperone are integrated specifically for simplifying workflows with agents: Agents work particularly well inside closed loops with verifiable rewards. Test mode reduces sampling for supported blocks to support small end-to-end checks, ideally using a persistent local GPU for shorter queue times on HPC clusters. This allows agents to adopt a fast build-test loop for complex pipelines (Fig 1**b**, right); efficiently testing and debugging the blocks integrate end-to-end as a fast verifiable signal. The same script can then scale to a full campaign, changed only through command-line overrides.

Additionally, blocks can run as separate steps, with each step’s returned DataFrame passed to the next and results gathered between jobs mimicking usual workflows (Fig. 1**d**, left). Alternatively, several blocks can be bundled into one call (Fig. 1**d**, right): Each chunk runs the multiple processes inside a single job, significantly cutting the scheduling overhead for multi-step workflows as well as improving throughput in cases with straggling processes.

## 3 *In silico* evaluation

To evaluate the impact of the Chaperone harness on agent-assisted protein design, we developed a *de novo* protein design evaluation suite spanning a diverse set of tasks and targets. Rigorously evaluating agent performance in this setting is challenging because successful *in silico* design depends on two intertwined capabilities: (1) the ability to execute and orchestrate complex, multistep tool-calling workflows and (2) the ability to reason about protein structure and make informed design decisions. To evaluate both capabilities, we provided agents with task-specific design specifications and the necessary molecular inputs, allowing them 12 hours and access to GPUs to autonomously develop, execute, evaluate, and iteratively refine their design strategies. At the end of each run, agents were required to submit 96 sequence-diverse candidate designs, which were independently evaluated using held-out scoring pipelines implemented through Chaperone as a benchmarking framework. These pipelines regenerated structures from the submitted sequences and assessed the resulting designs against predefined structural, geometric, and energetic criteria derived from real-world protein design tasks. We compared agent performance with and without access to the Chaperone tool-use harness, measuring the fraction of submitted designs satisfying all required criteria alongside computational resource consumption and token usage.

For our evaluation, baseline agents without the harness were allowed web access to install whichever tools they considered most suitable for accomplishing the design brief, while agents with the Chaperone harness had internet access disabled, as the necessary tools were already available. Evaluations were run using Harbor Controller [28], with agents given access to a maximum of five B4000 GPUs through a lightweight SLURM scheduler shim to track hourly usage and cost. For evaluations conducted with the harness, we provided a git-state-pruned copy of the Chaperone repository (to prevent leakage through the repository’s history), with relevant overlapping examples removed and containers pre-installed. GPU usage was similarly tracked through an environment shim. This comparison measures the practical benefit of the harness for end-to-end design under the evaluation conditions, although it does not disentangle workflow orchestration from structural reasoning and experimental activity. Additionally, evaluations were conducted on real, shared cluster resources with priority queuing to better reflect realistic computational workflow environments.

Across nine autonomous evaluations encompassing diverse protein design objectives and computational tools, GPT-5.6-sol agents with the Chaperone harness operating at high reasoning effort consumed 86% fewer total tokens and 64% fewer GPU-hours relative to baseline agents operating without the harness. Despite this substantial reduction in computational expenditure, the mean computational design pass rate increased from 6% to 12% (Fig. 1**f**). These findings indicate that improvements in computational efficiency need not come at the expense of design performance, and suggest that the burden of tool integration and workflow management may itself constitute a substantial bottleneck in autonomous protein design.

To further investigate how orchestration overhead scales with agent reasoning effort, we extended our evaluation across low, medium, and high reasoning settings for both configurations across the same nine tasks above. As expected, increasing reasoning effort was accompanied by greater token consumption, GPU utilization, and agent-issued actions in both settings. However, the magnitude of this increase differed between configurations. Baseline agents exhibited a median increase of 25 million tokens between successive reasoning-effort levels, whereas Chaperone-harnessed agents consumed fewer than 10 million tokens total - even at the highest reasoning setting (Fig. 1**g**). This divergence suggests that additional reasoning capacity in baseline agents is increasingly consumed by the implementation and maintenance of computational workflows, rather than necessarily translating into more extensive exploration of design strategies. In contrast, Chaperone’s standardized interfaces allow agents to construct, modify, and evaluate increasingly complex workflows without repeatedly regenerating the supporting scaffold. We additionally observed a substantial reduction in repeated job-status queries among Chaperone-harnessed agents, consistent with the availability of standardized job submission, automatic chunking, parallel execution, and failure recovery mechanisms that reduce the need for continuous intervention during long-running computations. Collectively, these observations suggest that the benefits of increased agent reasoning effort are strongly dependent on the computational environment in which the agent operates, with orchestration overhead potentially limiting the extent to which additional reasoning translates into productive scientific exploration.

## 4 Experimental results

### 4.1 Small molecule binder design

Designing high-affinity small-molecule binders remains challenging, with no single computational strategy generalizing effectively across targets. Differences in ligand geometry, interaction networks, and energetics often require extensive exploration of generative models, sequence design methods, and evaluation criteria to identify a suitable workflow. Previous efforts targeting cortisol involved iterative computational design followed by experimental affinity maturation, with Lee et al. ultimately achieving a binding affinity of 68 nM [29]. We therefore sought to investigate whether Chaperone could accelerate this process by allowing researchers and AI agents to collaboratively explore diverse design hypotheses, evaluate promising strategies through small-scale experiments, and selectively scale the optimal workflow.

We applied this approach to cortisol binder design, beginning with a sweep of 20 protein center-of-mass (COM) placements across four binder lengths, yielding 80 RFD3 runs to explore alternative pocket geometries and ligand contact surface strategies. From these, the researcher selected 10 promising backbone configurations for further sequence optimization, evaluating six customized LigandMPNN [7] amino-acid composition biases designed to promote hydrophobic packing around the ligand. The top three configurations were then subjected to iterative structure-prediction and refinement cycles, comparing six prediction-oracle combinations and six filtering strategies. Chaperone enabled the agent to execute and monitor 104 small-scale computational runs under designer guidance, progressively narrowing the design space before selecting a final workflow for large-scale generation (Fig. 2**b,c**).

**Figure 2.**
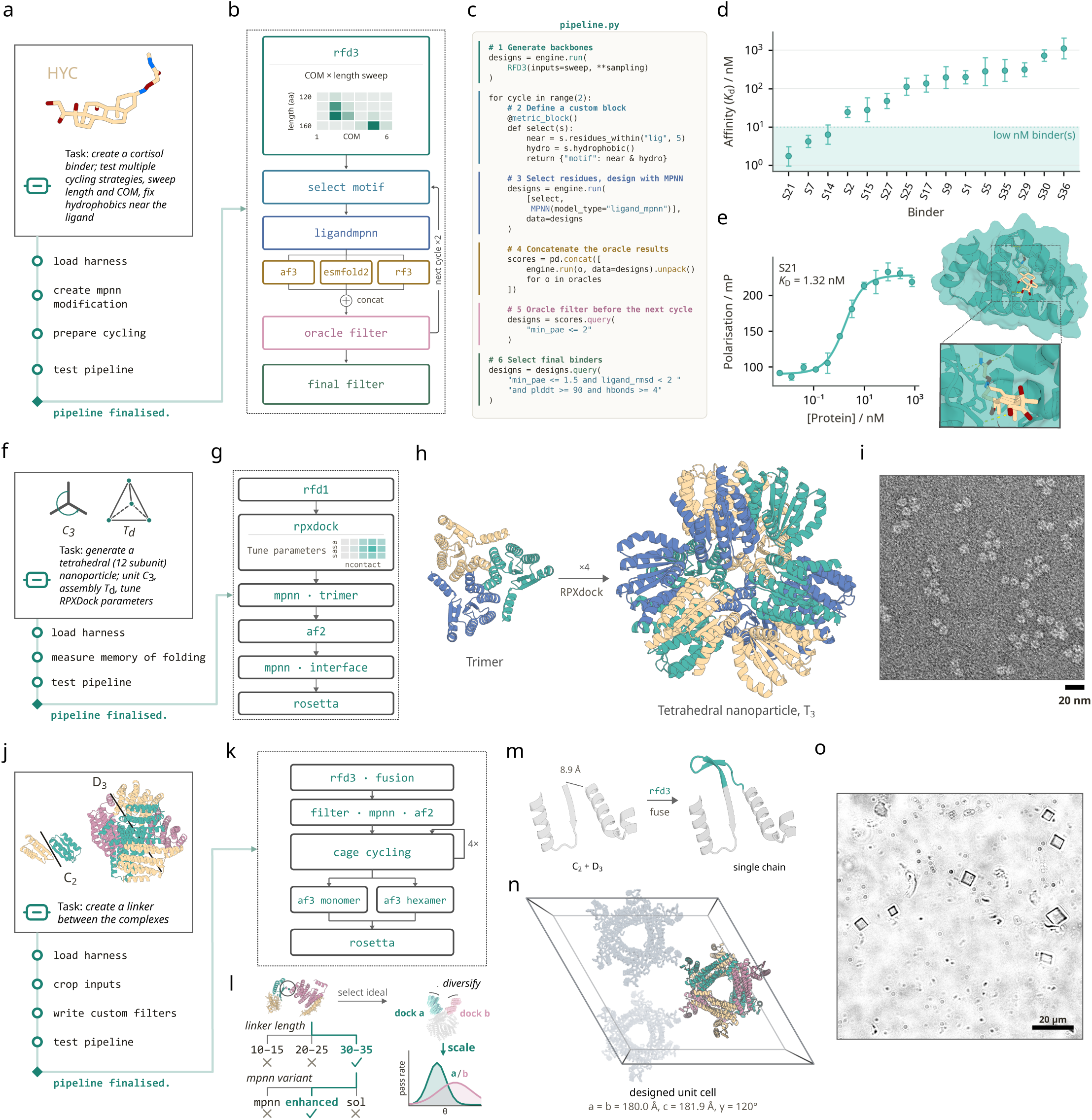
Experimental validation of multistep workflows with agent tool use. **a**, Small Molecule Protein Binder design with Chaperone: Agent is provided with the cortisol (HYC) ligand, as well as several multi-step workflow design strategies. **b**, Multi-step pipeline. RFD3 backbone generation with center of mass (COM) and length sweep, hydrophobic motif selection, ProteinMPNN sequence design with fixed motifs, and oracle cycling with AlphaFold3 (AF3), ESMFold2, and RoseTTAFold3 (RF3) applying a relaxed filter between cycles (min_pae < 2, hydrogen bonds ≥ 2). Designs after 2 cycles go through a final strict filter (min_pae < 1.5, ligand_rmsd < 1.0, pLDDT > 90). After sweeping several pipeline choices at pilot scale, the workflow then scales up. **c**, The complete pipeline script written by agent, separated by colors to its corresponding block on fig B, showcasing simplicity of concatenating mainstream tools alongside with a custom select motif block defined with the metric block decorator. **d**, Affinities (*K_D_*) of 15 designed binders with supported fluorescence-polarization fits and no quality-control flags, selected from 27 tested candidates; nine candidates with flagged fits and three with no clear response are omitted. The shaded band marks low-nanomolar binders. **e**, Fluorescence-polarization titration of best binder, with a fit of *K_D_* = 1.32 nM. Design model of the binder bound to HYC, showing the binding site with key hydrogen bonds highlighted. **f**, Nanoparticle design with Chaperone: Agent is provided with the desired symmetry groups of the design, as well as details of how to carry out the task. **g**, The devised pipeline consists: RFD1 backbone generation, RPXdock docking, ProteinMPNN sequence design for the trimer, AF2 structure prediction, ProteinMPNN interface design and Rosetta scoring. **h**, Design model of the 12-chain tetrahedral nanoparticle. **i**, Negatively stained electron micrograph of the nanoparticle, revealing assemblies of the expected size. Scale bar = 20 nm. **j**, Linker-design with Chaperone: Agent is tasked to create a linker between two symmetric complexes. Chaperone crops the inputs, writes custom filters, and devises the design pipeline shown in k. **k**, The devised pipeline. RFD3 builds a single-chain fusion. Backbone filters, ProteinMPNN and AF2 run as one block. CaGE cycling is repeated four times. AF3 monomer and hexamer predictions run in parallel and merge into Rosetta scoring. **l**, Illustration of pilot sweep and input diversification. At pilot scale (∼100 designs) the agent swept linker length and the MPNN variant (ProteinMPNN [2], SolubleMPNN [19] and EnhancedMPNN [32].) The ideal length and sequence variant were selected to fix the base pipeline, and subsequent design parameters - multiple docks and arbitrary fixed C2/D3 relative complex orientation - were assessed during scaling. **m**, Zoom on the linker of design H11. Left, the two prepositioned complexes, with the 8.9 Å gap between the termini to be joined (dashed). Right, RFD3 fuses them into a single chain; the designed 17-residue linker, a *β*-hairpin, is shown in teal. **n**, Unit cell of the H11 crystal design (space group R 3 2:H; a = b = 180.0 Å, c = 181.9 Å, *γ* = 120^◦^) viewed down c. One complete assembly of six chains is coloured by chain; the other two assemblies in the cell are grey. **o**, Brightfield microscopy of H11 crystals grown in an elution buffer.

This exploration went beyond conventional parameter optimization, allowing us to modify the customized pipeline blocks as new hypotheses emerged, without repeatedly implementing the infrastructure needed to connect individual tools. Within two weeks, we progressed from initial design hypotheses to sequences ready for experimental testing. Of 27 candidates tested by fluorescence polarization in the first experimental round, 15 yielded supported affinity fits without quality-control flags, nine yielded flagged fits, and three showed no clear response. The best binder had an affinity of 1.32 nM without any affinity maturation. This result highlights Chaperone’s utility in making extensive methodological exploration practical within a short development cycle, allowing researchers to spend less effort managing computational workflows and more time investigating the design strategies most likely to succeed (Fig. 2**d,e**).

### 4.2 Nanoparticle design

Designing *de novo* self-assembling protein nanoparticles has been historically challenging and often consists of many different steps. A recent approach described the design of *de novo* protein nanoparticles [30] used the following steps: (a) design *de novo* symmetric oligomers around a motif with RFdiffusion [5], (b) dock those oligomers into the desired geometry with RPXdock [31], (c) design the sequence of the entire nanoparticle with ProteinMPNN, and (d) use AlphaFold [10] and Rosetta [13] to score and select constructs. Here, we used a similar approach to design protein nanoparticles with Chaperone as a demanding test case for its ability to incorporate uncommon software, like RPXdock, into a standard pipeline (Fig. 2**g**).

We aimed to design tetrahedral nanoparticles containing a trimeric coiled-coil motif. By providing the agent with a design outline for the approach previously described, it easily generated a small library of tetrahedral nanoparticles from which we could filter and select 15 designs for experimental validation. Of these, one successfully assembled into a protein nanoparticle, a success rate of 6.7% (Fig. 2**h,i**). This result demonstrates the ability of Chaperone to translate an established design approach into a streamlined pipeline that produces experimentally validated proteins.

### 4.3 Protein fusion

Rigidity is a central requirement in crystal design, since the lattice only repeats if the components hold the orientation they were given. Designing a rigid linker (Fig. 2**m**) that scaffolds two symmetric motifs and keeps their relative geometry fixed is therefore the step the rest of the crystal depends on, and the length that achieves it is not known in advance, although most tools require it to be fixed a priori.

We applied this approach to the fusion of C2- and D3-symmetric components, where the agent used RFD3 to build single-chain fusions from prepositioned motifs and customized filters to discard backbones that clash with symmetry mates or connect the motifs too weakly. The prepositioned motifs were generated by a separate rigid-body placement and scoring pipeline developed in our laboratory for *de novo* protein crystal design (manuscript in preparation), and are used here as fixed inputs. A base pipeline is devised based on the desired constraints (Fig. 2**k**), and agents undertake pilot experiments to determine ideal length, the ideal ProteinMPNN variant, and whether to adopt the cycling strategy (Fig. 2**l**). We then selected the ideal parameters and scaled backbone generation to ∼10x the pilot, diversifying the inputs across ∼4,500 placements instead of one. We tested seven lengths per placement, with the window centre taken from the pilot for 4,355 placements (97.9%) and from a recorded fallback for the remaining 94. Both rounds used the same pipeline, with the additional load automatically dispatched by the execution engine.

Designs were refined through templated structure prediction and sequence redesign cycles using CaGE [18], with specified residues fixed to preserve crystallization interfaces. Classifier-free guidance was annealed over four stages, and a cycled backbone was kept only if its fixed residues stayed within 1.0 Å backbone RMSD of the input. Designs were then evaluated with hexamer AlphaFold3 predictions and researcher-provided analysis scripts. This example demonstrates Chaperone’s ability to settle a design parameter in a disposable pilot and carry the same workflow into production.

## 5 Conclusions

In this work, we showed that generalized design across a range of tasks with freely available protein design tools can be standardised under simple principles. The design of *blocks* allows for a simple interface that allows rapid integration across a wide array of biomolecular tools. The harness design of Chaperone incorporates multiple optimizations to biomolecular design workflows such as incorporating verification loops for complex pipelines, bundled multi-step jobs with customised augmentation and manipulation of structures at scale. We demonstrate the harness improves agent tool use on real design workflows, owing to simplified orchestration, efficient context management, improved output formatting and job polling, as well as pre-built pipeline patterns. We found the workflows of harnessed agents are more monitorable, owing to their reduced output space, enabling reliable and manageable scaling to several agents or complex tasks.

We demonstrated design with Chaperone across multiple wet-lab validated tasks: We first demonstrated the design of high affinity (low nanomolar) cortisol binders with RFdiffusion3 as a generation oracle, scaled with customised hydrophobic sequence design tailored to the design task. Chaperone enables such a complex workflow to be fully tested and encapsulated by a few dozen lines of code, rather than thousands. We further demonstrated how non-standard design tasks—requiring tools external to the conventional generate-predict pipeline—are particularly well suited for the Chaperone framework: we demonstrate the successful design of a 12-subunit nanoparticle, using autonomous RPXDock parameter tuning. Finally, we demonstrate the task of linker design can be effectively carried out; by tuning key design parameters such as length and pipeline choices such as sequence fitting method. These workflows yielded observed crystals with brightfield microscopy.

We note that rigorously evaluating tool use is compute intensive; evaluations mix stochasticity within the tool calls themselves with the agents and tool use choices. We observed task-by-task variability, for example in protein binder design. As a result, we chose to more deeply evaluate GPT-5.6 Sol with the resources available. We leave a large-scale evaluation of agent tool use across model families and harnesses as directions for future work.

Overall, Chaperone supports a wide range of tasks and enables scientists to tackle ambitious long-horizon design problems with more reliable agents. Additionally, Chaperone can readily be used in a number of additional contexts not explored here; including retrospective analysis, model benchmarking and development. As tools increase in generation and prediction quality, we expect the necessity to effectively orchestrate and couple tools together and to orchestrate long-horizon agentic tasks to become increasingly prominent. To this end, we make Chaperone freely available to the community on GitHub: https://github.com/RosettaCommons/chaperone.

## Author contributions

Scientific design: JB, JC, DB. Infrastructure design: JB, AH, JC. Design and validation of nanoparticles: CMH. Design and wet lab validation of small molecule binders: JC, SS. Design and wet lab validation of protein fusion: WD. C2-symmetric component: ZL. D3-symmetric component: HP. Agent benchmarking: JC, JB. Software and development: JB, JC, AH, SS, JK, AM, RM, RB, MB. Writing and review: All authors.

## Acknowledgements

We thank the IPD staff: Luki Goldschmidt and Patrick Vecchiato for their instrumental help in aiding with the development of HPC orchestration engines. We thank Andrew S Rosen for insightful discussions related to software design. We thank Ian Anderson for his valuable feedback for development. We thank Joy Jiao from OpenAI for their support in the form of Codex credits for the IPD. Finally, we thank Jonah Cool and Bailey Bova at Anthropic for their support in the form of credits for the IPD in the early development of this project.

This work was supported by the National Institute of Allergy and Infectious Diseases (U19AI181881 to N.P.K. and D.B.); the Advanced Research Projects Agency for Health (ARPA-H; APECx Program Award No. 1AY1AX000036-01 to N.P.K. and D.B.); the National Science Foundation Graduate Research Fellowships Program Award (GR068505 to J.C. and DGE-2140004 to C.M.H.); the Grantham Foundation (J.B., D.B.); the IPD Enzyme Design Fund (J.C.); the Howard Hughes Medical Institute (D.B.); The Audacious Project at the Institute for Protein Design; and the Open Philanthropy Project Improving Protein Design Fund.

## Supplementary information

### S1#Design cases for agent evaluation

In this section, we describe the design cases for each task in the evaluation described in Figure 1. Our evaluation spans a breadth of protein design tasks. Every task follows a similar architecture: (1) a prompt which specifies the design task, input files, and criteria, (2) a held-out scoring function which measures the success of designs that is not visible to agents completing the benchmark, (3) a submission contract which is strictly enforced for designs to be scored. All submission contracts enforce the following requirements: submission must be delivered in a flat CSV with exactly 96 designs, protein sequences must be uppercase amino acid letters only, and pairwise sequence identity (Needleman–Wunsch [33]) between designs must be ≤ 0.7. For valid submissions, scoring outputs both a primary score satisfying all hard criteria, and a second ‘soft_pass’ score that reports designs passing both hard and any specified additional soft criteria.

#### S1.1#Protease

Model inputs and criteria: A reference theozyme comprising six catalytic residues (H293, E294, H297, E316, Y377, and E378) arranged around zinc oxide, supporting motifs, and the fibrin A*α* substrate peptide GPRVVERHQS are supplied for *de novo* zinc metalloprotease design. Catalytic-residue all-atom RMSD must be ≤1.5 Å. Submissions must include a motif_index_map mapping reference residues to distinct sequence positions with matching amino acid identities. Additional scorer operations: AF3 predicts the enzyme–substrate complex. The scorer evaluates motif fidelity and additionally requires whole-complex pLDDT >0.8 and minimum protein–substrate interface PAE ≤10 Å.

#### S1.2#Symmetric assembly

Model inputs and criteria: The task specifies a compact C3-symmetric homotrimer. An illustrative reference protomer and generation configuration suggest a sequence length of 60–90 residues but do not affect scoring. Public hard criteria require intersubunit rotation deviation ≤5^◦^ from 120^◦^, aligned protomer RMSD ≤1 Å, representative A–B interface contact molecular surface ≥600, chain-A loop fraction <0.35, chain-A C*α* radius of gyration ≤14 Å, zero backbone clashes, ipTM ≥0.85, and pLDDT ≥85. Interface ΔΔG <−35 is a soft criterion. Additional scorer operations: Each sequence is replicated into three identical chains and the assembly is predicted using AF3. All disclosed criteria are evaluated on this structure; no additional thresholds are imposed.

#### S1.3#Peptide binder

Model inputs and criteria: The EIN2 site-1 peptide TVGSDGPPSFRSLSG is supplied in a specified conformation as chain B, with all residues designated as hotspots. The binder constitutes chain A, with a suggested length of 60–150 residues. Public hard criteria require aligned backbone RMSD ≤1.5 Å over peptide residues B9–B15, binder loop fraction <0.35, binder–peptide contact molecular surface ≥300, zero backbone clashes, and ipSAE [34] ≥0.30. Interface ΔΔG ≤−15 is a soft criterion. Additional scorer operations: AF3 predicts the complex without peptide templating to assess recovery of the supplied epitope conformation. The scorer evaluates the public criteria and additionally requires minimum interface PAE ≤1.5 Å and chain-A pLDDT ≥0.85.

#### S1.4#RNA binder

Model inputs and criteria: The target is the 20-nucleotide stem-loop 5^′^-GAGGACCUAGAUCACCCCUC-3^′^ from PDB 1EC6. Inputs include DSSR [35] stem annotations (G1–C20, A2–U19, G3–C18, G4–C17, and A5·C16) and an RFD3 configuration specifying loop-face hotspots and hydrogen-bond conditioning. Public criteria require ≥8 protein–RNA hydrogen bonds, ≥2 minor-edge bonding interactions, ≥12 protein–RNA contacts, contact molecular surface ≥250, ≥2 base-stacking interactions, pLDDT ≥0.85, zero backbone clashes, and base-paired stem backbone RMSD ≤1 Å. Additional scorer operations: AF3 predicts the complex, DSSR/SNAP annotates nucleotide interactions, and backbone alignment determines stem RMSD. The scorer additionally requires a minimum interface PAE ≤2 Å.

#### S1.5#Protein–protein binding: PD-L1

Model inputs and criteria: PD-L1 residues 18–132 from PDB 4ZQK are supplied as trimmed, relabeled chain B, with hotspots B56, B66, B115, B121, B122, B123, B124, and B125. Public criteria specify binder chain-A loop fraction <0.3, interface ΔΔG ≤−25, and symmetric interface PAE ≤7 Å. Additional scorer operations: The scorer evaluates these criteria and additionally requires minimum interface PAE ≤1.25 Å.

#### S1.6#DNA binder

Model inputs and criteria: The task follows the DNA-binding design setting of Sehgal et al. [36]. A B-form Dickerson–Drew dodecamer duplex (CGCGAATTCGCG) is supplied as two chains, with sequence-specific recognition directed to the major groove. Public criteria require ≥3 major-groove hydrogen bonds, zero protein backbone clashes, interface contact molecular surface ≥400, ≥8 amino acid–nucleotide hydrogen bonds, ≥10 amino acid–nucleotide contacts, and nucleotide-backbone-aligned DNA RMSD ≤2 Å. Additional scorer operations: An independent held-out structural oracle evaluates these criteria and additionally requires interface PAE ≤5 Å, worst cross-chain ipSAE ≥0.3 across both DNA chains, and protein pLDDT ≥80.

#### S1.7#Ligand binder

Model inputs and criteria: Linker-modified biotin is specified by its SMILES, atom names, and regional exposure requirements. Public criteria require solvent-accessible surface area ≥0.1 for at least one linker atom (C14, C15, or C16), ligand-core contact molecular surface ≥120, ≥3 protein–ligand hydrogen bonds, and ≥1 aromatic residue within 6 Å of the ligand. Additional scorer operations: AF3 predicts the complex. The scorer evaluates the public criteria, additionally requires minimum interface PAE ≤1.5 Å, and increases the ligand-core contact molecular surface threshold to ≥140.

#### S1.8#Antibody CDR redesign

Model inputs and criteria: The antibody light chain, heavy chain, and lysozyme antigen from PDB 1VFB are supplied with six designated CDR regions. Five CDR lengths are fixed; H3 may contain 5–20 residues. Public hard criteria require fixed-framework backbone RMSD ≤1.5 Å, antibody–antigen contact molecular surface ≥300, and zero backbone clashes. Soft criteria require no NXS/T sequons, ≤1 unpaired cysteine, net charge between −5 and +5, and no H3 run exceeding four hydrophobic residues (ACFILMVWY). The submission contract requires framework sequence preservation and measures pairwise sequence identity ≤0.7 only over redesigned CDRs. Additional scorer operations: AF3 predicts the complex. The scorer evaluates the public criteria and additionally requires minimum antibody–antigen interface PAE ≤8 Å, chain-pair-normalized interface ipSAE ≥0.05, and whole-complex pLDDT ≥0.8.

#### S1.9#Glycosidase enzyme

Model inputs and criteria: A theozyme containing Y77, E86, Y88, R122, Q136, and E177 around 4-methylumbelliferyl *β*-glycoside is supplied. Public hard criteria require catalytic side-chain all-atom RMSD ≤1 Å, catalytic glutamate-dyad distance of 5.9–8.9 Å, motif-aligned substrate RMSD ≤2 Å, mean substrate solvent accessibility ≤0.15, and protein–substrate contact molecular surface ≥200. ≥3 protein–substrate hydrogen bonds is a soft criterion. Submissions must include a catalytic-residue motif_index_map. Additional scorer operations: AF3 predicts the complex. The scorer evaluates the public criteria and additionally requires whole-complex pLDDT ≥85 and ligand pLDDT ≥70.

### S2#Wet lab validation of nanoparticles

#### S2.1#Protein purification

DNA fragments for expression and screening of protein designs (IPDblocks) were generated from oligonucleotide pools using the IPDassembly process, similar to previous reports. Briefly, DNA sequences encoding designed proteins were bioinformatically fragmented, and each fragment was appended with BsaI restriction sites and design-specific PCR priming sites. These sequences were synthesized within a 350 nucleotide oligonucleotide pool at Twist Bioscience. Two nested PCR stages were conducted to dial-out design-specific fragments within each well of a microwell plate. DNA fragments composing each design were individually combined with LM1371 and assembled via a Golden Gate reaction [37]. Genes were then transformed into BL21 competent *E. coli* cells (NEB). Constructs were screened at 1 mL scale where cells were grown in 96-deep well format in autoinduction media [38] (Terrific Broth II media supplemented with 50×5052, 50×M salts, 20 mM MgSO_4_, and a trace metal mix) at 37^◦^C for 24 hours at 1350 revolutions per minute (rpm) in a Heidolph shaker. Cells were then spun down at 4000 g for 30 minutes and resuspended in lysis buffer (25 mM Tris pH 8, 250 mM NaCl, 20 mM Imidazole, 200 mM L-Arginine) with the addition of bovine pancreatic DNase I (Sigma-Aldrich) and protease inhibitors (Thermo Scientific). Resuspended cells were then sonicated with a 24-prong sonicator. Lysed cells were centrifuged for 30 minutes at 4000 g and clarified lysate was subsequently poured over equilibrated Ni-NTA resin (QIAGEN) in a gravity column. The columns were washed with 10 column volumes of wash buffer (25 mM Tris pH 8, 250 mM NaCl, 20 mM Imidazole), then mixed with and incubated in elution buffer (25 mM Tris pH 8, 300 mM NaCl, 500 mM Imidazole) for 10 minutes prior to being eluted. Eluant was concentrated to 100 *μ*L and further purified using Superdex 200 5/150 columns on an HPLC (Agilent 1260 Infinity II LC System) and buffer exchanged into a sizing buffer (25 mM Tris pH 8, 500 mM NaCl, 100 mM L-Arginine) before selecting fractions for confirmed assembly via negative stain electron microscopy.

#### S2.2#Negative stain electron microscopy

300-mesh carbon coated grids (Electron Microscopy Sciences) were glow discharged for 30 seconds prior to application of protein samples (3 *μ*L, 0.1 mg/mL) and blotting three times with 2% (w/v) uranyl formate. Micrographs were acquired using a 120 kV Talos L120C transmission electron microscope (Thermo Scientific) with a BM-Ceta camera.

### S3#Wet lab validation of protein fusion

#### S3.1#Protein expression and purification

Selected designs were expressed from a customized pET29b-derived vector with a C-terminal His-tag (LM627, Addgene 191551). Plasmids were transformed into chemically competent *E. coli* BL21(DE3) cells. Single transformants were used to inoculate 1 mL LB pre-cultures supplemented with 50 *μ*g/mL kanamycin, which were grown at 37 ^◦^C for 16 h. Pre-cultures were used to inoculate expression cultures in autoinduction medium [38] (Terrific Broth II supplemented with 25× M salts, 25× 5052, 2 mM MgSO_4_, a trace metal mix and 50 *μ*g/mL kanamycin), distributed as 1 mL per well in 2 mL 96-deep-well plates, for a total of 4 mL per design. Expression cultures were grown at 37 ^◦^C for 20 h without a temperature shift. Cells were harvested by centrifugation at 4,000 × g for 15 min at 16 ^◦^C, and the pellets were frozen at −80 ^◦^C overnight.

Lysis buffer was prepared by combining 5 mL BugBuster, approximately 100 mg lysozyme, 1 mL bovine pancreatic DNase I, 50 *μ*L of 1 M MgSO_4_ and one protease inhibitor tablet, topped up to 50 mL with wash buffer (50 mM Tris pH 8, 150 mM NaCl, 20 mM imidazole pH 8). Frozen pellets were resuspended in lysis buffer at 100 *μ*L per 1 mL of expression culture and lysed by shaking at room temperature for 1 h; no sonication was used. The lysate was then incubated with Ni-NTA beads for 15 min at room temperature. The beads were washed once with wash buffer and once with high-salt wash buffer (50 mM Tris pH 8, 1 M NaCl, 20 mM imidazole pH 8), and bound protein was eluted in elution buffer (50 mM Tris pH 8, 150 mM NaCl, 400 mM imidazole pH 8).

#### S3.2#Bright-field imaging

Purified protein was transferred to a black 96-well glass-bottom plate with #1.5 high-performance cover glass (0.17 ± 0.005 mm; Cellvis, P96-1.5H-N) and imaged on an IN Cell Analyzer 2500 HS (Cytiva, formerly GE Healthcare) using bright-field illumination at 60× magnification.

### S4#Wet lab validation of cortisol binder

#### S4.1#Materials

For binding experiments, Hydrocortisone (Sigma Aldrich H4001, synonym: cortisol) was used as the design target throughout the computation pipeline and as a competitor. TAMRA (5-Carboxytetramethylrhodamine, Thermofisher C6121) was used as a negative control. TAMRA-PEG3-cortisol [29] was a custom synthesized ligand used for fluorescence activated cell sorting and fluorescence polarization experiments.

#### S4.2#Yeast surface display screening for cortisol binders

Designed protein sequences were reverse-translated using DNAWorks [39] and codon-optimized for *Saccharomyces cerevisiae*, excluding common restriction enzyme recognition sites. Each coding sequence was divided into overlapping 5^′^ and 3^′^ fragments, and designs were grouped by length to minimize length-dependent amplification bias. The corresponding oligonucleotides were synthesized as a 300-nt oligonucleotide pool (Twist Bioscience). The 5^′^ and 3^′^ fragments were amplified separately by quantitative PCR (qPCR) using KAPA HiFi HotStart Uracil+ ReadyMix. Inner primer-derived adapter sequences were removed by sequential treatment with USER enzyme and the NEBNext End Repair Module, and corresponding fragments were assembled into full-length inserts by overlap-extension qPCR using KAPA HiFi HotStart ReadyMix. The assembled insert pool (5 *μ*g) was combined with linearized pETCON3 vector (25 *μ*g) and introduced into electrocompetent *Saccharomyces cerevisiae* EBY100 cells following a previously published protocol [40]. Transformants were cultured in complete synthetic medium lacking tryptophan and uracil and supplemented with 2% glucose (CTUG) at 30 ^◦^C for 48-72 h, until an OD_600_ of 4-5 was reached. Surface display was subsequently induced in SGCAA medium at 30 ^◦^C for 18 h.

**Figure S1.**
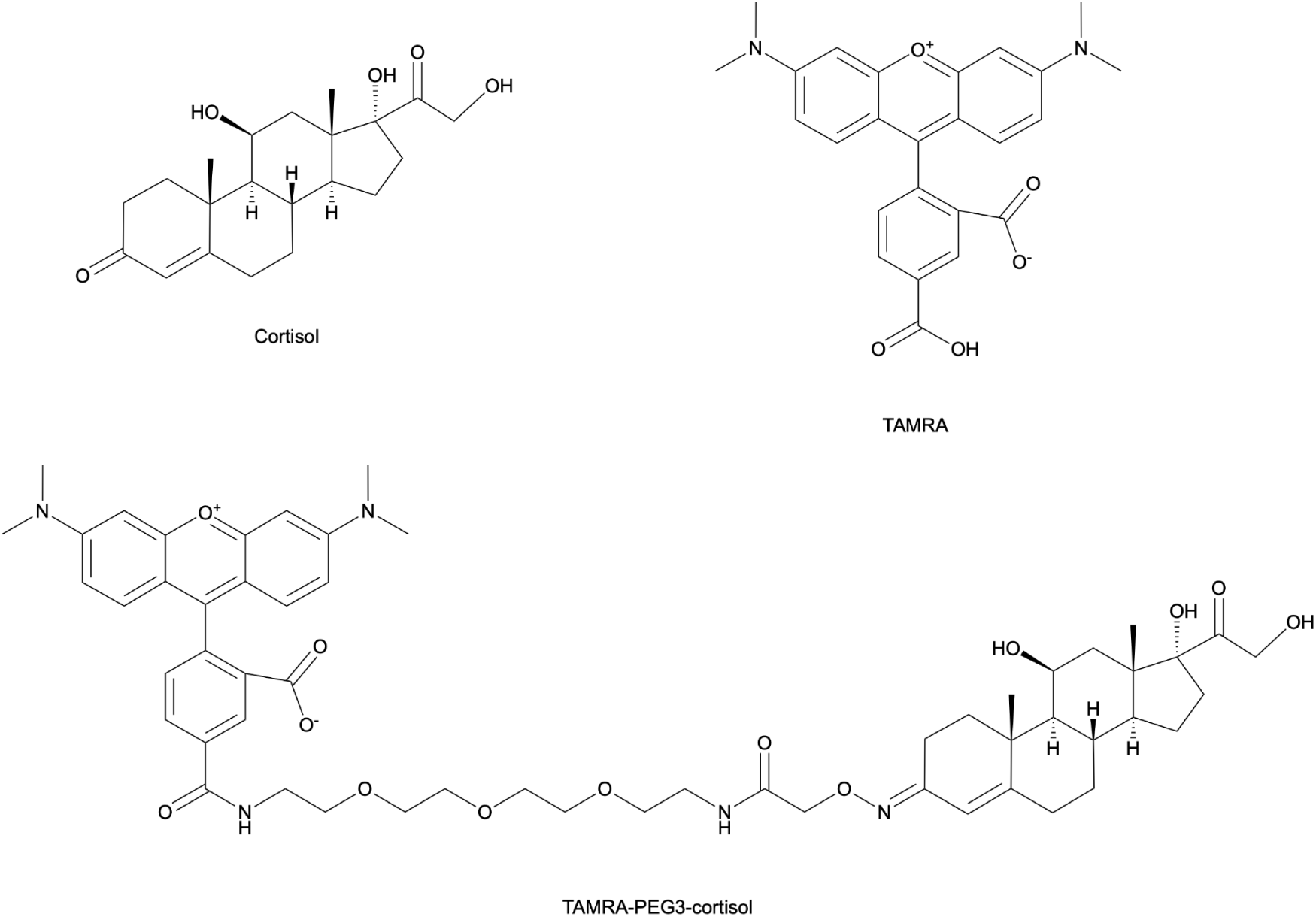
Chemical structures of cortisol, TAMRA, and TAMRA-PEG3-cortisol.

All staining and ligand-binding incubations were performed in phosphate-buffered saline (PBS, 137 mM NaCl, 2.7 mM KCl, 10 mM Na_2_HPO_4_, 1.8 mM KH_2_PO_4_, pH 7.4) supplemented with 0.1% (w/v) bovine serum albumin (PBSF). Fluorescence-activated cell sorting (FACS) was performed on a Sony SH800S equipped with 488- and 561-nm lasers, using a 100-*μ*m sorting chip and PBS as the sheath fluid. An initial expression sort was performed to enrich the library for designs displaying detectable levels of the C-terminal c-Myc tag. 20 million induced cells were washed twice with 1 mL PBS, resuspended in PBSF containing FITC-conjugated anti-c-Myc antibody (Immunology Consultants Laboratory, CMYC-45F) at a 1:50 (v/v) dilution, and incubated for 30 min at 4 ^◦^C. Following two washes with PBSF, c-Myc-positive cells were collected by FACS, expanded in CTUG medium to an OD_600_ of 4-5, and re-induced in SGCAA medium for 18 h.

For binding enrichment, induced cells were stained with FITC-conjugated anti-c-Myc antibody as described above, washed twice with PBSF, and divided into experimental and control aliquots. Experimental aliquots were incubated with 1 *μ*M TAMRA-PEG3-cortisol, whereas control aliquots were incubated with 1 *μ*M TAMRA alone, both for 60 min at 4 ^◦^C. Cells were subsequently washed twice with PBSF. From the TAMRA-PEG3-cortisol-treated population, FITC-positive cells exhibiting TAMRA fluorescence above the TAMRA-only control were collected, with TAMRA fluorescence measured in the PE detection channel. Two rounds of binding enrichment were performed. For library titrations, cells were incubated with TAMRA-PEG3-cortisol at seven concentrations: 0, 320 pM, 1.6 nM, 8 nM, 40 nM, 200 nM, and 1 *μ*M and sorted using the same gate across all concentrations. Binding volumes were scaled with ligand concentration to minimize ligand depletion.

Plasmid DNA was isolated from sorted yeast populations using the Zymoprep Yeast Plasmid Miniprep II kit (Zymo Research), and the design-encoding regions were amplified by PCR using indexed primers. Amplicon libraries were sequenced on an Illumina NextSeq 1000 using 2 × 300-bp paired-end reads. Paired-end reads were assembled and matched to the reference design sequences using a custom analysis pipeline [41] yielding per-design read counts for each sorted population.

For titration sorts, concentration-dependent enrichment was used to estimate SC_50_ as a proxy for apparent dissociation constants (*K_D_*) for binding following a previously published analysis procedure [41] . Designs with estimated apparent *K_D_* values ≤1 *μ*M were classified as candidate binders.

#### S4.3#Protein expression and purification

DNA fragments encoding candidate binders were ordered as Twist Gene Fragments (Twist Bioscience) with flanking sequences for Golden Gate cloning [37] using BsaI-HFv2 (New England Biolabs R3733L). Each fragment was individually assembled into the LM670 expression vector (Addgene plasmid 191552), and the assembly products were transformed into chemically competent *Escherichia coli* BL21 cells (Intact Genomics 1054-81). Each design was expressed in four parallel 1-mL cultures distributed across four 96-deep-well plates (Fisher Scientific PDW20CSIND). Cultures were grown in autoinduction medium [38] consisting of Terrific Broth II (Fisher Scientific MP113046052) supplemented with 1x 5052 solution and 1 mM MgSO_4_ at 37 ^◦^C for 4 h, followed by 20 ^◦^C for 24 h, with shaking at 1,350 rpm on a Heidolph orbital shaker. Cells were harvested by centrifugation at 4,000 × g for 15 min, and pellets were stored at -20 ^◦^C overnight. Pellets were resuspended in lysis buffer (100 mM Tris, pH 8.0, 300 mM NaCl, and 25 mM imidazole) supplemented with BugBuster 10x Protein Extraction Reagent diluted to 1x (MilliporeSigma 70921-5) bovine pancreatic DNase I (Sigma-Aldrich DN255G), and protease inhibitor tablet (Thermo Fisher A32965). Following resuspension, the four samples corresponding to each design were pooled. Lysates were clarified by centrifugation at 4,000 × g for 30 min. Each clarified lysate was applied to 100 *μ*L of pre-equilibrated Ni-NTA resin (QIAGEN 30250-A) in a 96-well plate with 25-*μ*m frits (Agilent 200953-100). The resin was washed with 10 column volumes of wash buffer (50 mM Tris, pH 8.0, 200 mM NaCl, and 25 mM imidazole). Proteins were eluted by incubating the resin with elution buffer (50 mM Tris, pH 8.0, 200 mM NaCl, and 250 mM imidazole) for 5 min, followed by centrifugation at 500 x g for 2 min. Eluates were filtered through a 0.2-*μ*m 96-well filter plate (Cytiva 8586) and further purified by size-exclusion chromatography (SEC) using a Superdex 75 Increase 5/150 GL column (Cytiva 29148722) on an Agilent 1260 Infinity II LC system. SEC was performed in phosphate-buffered saline (PBS, 137 mM NaCl, 2.7 mM KCl, 10 mM Na_2_HPO_4_, and 1.8 mM KH_2_PO_4_, pH 7.4).

#### S4.4#Fluorescence polarization

Purified proteins were used directly to prepare dilution series in PBS. Stock concentrations were determined from absorbance measurements at 280 nm using a NanoDrop spectrophotometer. Fluorescence polarization (FP) assays were performed in 96-well plates (Corning 3686). Each well initially contained 50 *μ*L PBS, and 25 *μ*L purified protein stock was added to the first well of each dilution series. Following mixing, 25 *μ*L was transferred sequentially between wells to generate a 12-point, threefold serial dilution. Subsequently, 50 *μ*L of 2 nM TAMRA-PEG3-cortisol was added to each well, yielding a final assay volume of 100 *μ*L and a ligand concentration of 1 nM. The initial screen used one dilution series per protein; the four selected proteins were subsequently measured using three technical replicate dilution series.

FP was measured using an Agilent BioTek Synergy Neo2 plate reader with excitation at 530 nm (25 nm bandwidth) and emission at 590 nm (20 nm bandwidth) using a red filter cube (Agilent 1035062). Measurements were collected using top-read optics, a gain setting of 120 and a read height of 7.5 mm, with ten instrument measurements per data point. The recorded reader temperature was approximately 24^◦^C. Parallel and perpendicular fluorescence intensities and polarization values in millipolarization units (mP) were exported for analysis.

For competition measurements of all candidate binders, 10 *μ*L of 1 mM unlabelled ligand was added to each well following the initial FP measurement. This increased the total volume to 110 *μ*L, yielding final concentrations of 90.9 *μ*M unlabelled ligand and 0.909 nM TAMRA-labelled ligand, with a corresponding dilution of protein concentrations by a factor of 100/110. The same wells were measured shortly after competitor addition using identical reader settings.

Apparent dissociation constants were determined by globally fitting triplicate fluorescence anisotropy titrations to the quadratic 1:1 binding model below accounting for ligand depletion [42]. The total labelled-ligand concentration was fixed at 1 nM.

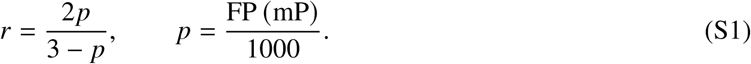

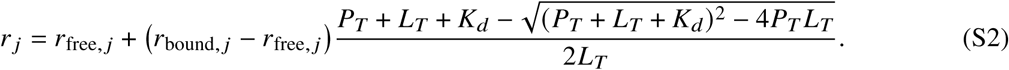

Here, *p* is fractional fluorescence polarization and *r* is fluorescence anisotropy. The index *j* denotes a technical replicate; *r*_free,_ *_j_* and *r*_bound,_ *_j_* are its fitted free- and bound-ligand anisotropies. *P_T_* and *L_T_* are the total protein and labelled-ligand concentrations, respectively, and *K_d_* is the apparent dissociation constant shared across replicates. Concentrations and *K_d_* are expressed in nM.

## Notes

### Competing Interest Statement

The authors have declared no competing interest.

